# USP7 as part of non-canonical PRC1.1 is a druggable target in leukemia

**DOI:** 10.1101/221093

**Authors:** Henny Maat, Jennifer Jaques, Aida Rodríguez López, Shanna M. Hogeling, Marcel P. de Vries, Chantal Gravesteijn, Annet Z. Brouwers-Vos, Nisha van der Meer, Gerwin Huls, Edo Vellenga, Vincent van den Boom, Jan Jacob Schuringa

## Abstract

Acute myeloid leukemia (AML) is a highly heterogeneous disease in which genetic and epigenetic changes disturb regulatory mechanisms controlling stem cell fate and maintenance. AML still remains difficult to treat, in particular in poor risk AML patients carrying TP53 mutations. Here, we identify the deubiquitinase USP7 as an integral member of non-canonical PRC1.1 and show that targeting of USP7 provides an alternative therapeutic approach for AML. USP7 inhibitors effectively induced apoptosis in (primary) AML cells, also independent of the USP7-MDM2-TP53 axis, whereby survival of both the cycling as well as quiescent populations was affected. MLL-AF9-induced leukemia was significantly delayed *in vivo* in human leukemia xenografts. We previously showed that non-canonical PRC1.1 is critically important for leukemic stem cell self-renewal, and that genetic knockdown of the PRC1.1 chromatin binding component KDM2B abrogated leukemia development *in vitro* and *in vivo* [1]. Here, by performing KDM2B interactome studies in TP53mut cells we identify that USP7 is an essential component of PRC1.1 and is required for its stability and function. USP7 inhibition results in disassembly of the PRC1.1 complex and consequently loss of binding to its target loci. Loss of PRC1.1 binding coincided with reduced H2AK119ub and H3K27ac levels and diminished gene transcription, whereas H3K4me3 levels remained unaffected. Our studies highlight the diverse functions of USP7 and link it to Polycomb-mediated epigenetic control. USP7 inhibition provides an efficient therapeutic approach for AML, also in the most aggressive subtypes with mutations in TP53.

**Key points:**

1. USP7 is a therapeutic target in leukemia, including poor risk TP53mut AML.
2. USP7 is an essential component of non-canonical PRC1.1 and is required for its stability and function.

## INTRODUCTION

Patients with AML often have a poor prognosis despite treatment with intensive chemotherapy and allogeneic stem cell transplantation. Dependent on risk category overall survival for adult AML patients varies between 10%-60% [2]. AML is a disease of the elderly accounting for 75% of the cases in patients >60 years of age which have a particularly poor outcome [3, 4]. These older patients usually have karyotypes associated with unfavorable risk and also TP53 mutations are more frequently seen in patients above 60, and this patient group generally does not respond well to standard chemotherapy [4–6]. Therefore, alternative therapies need to be developed to achieve more effective treatment of AML patients.

A recurrent challenge in AML treatment is the notion that standard-of-care chemotherapeutic approaches do not effectively target quiescent leukemic stem cell (LSC) populations, and as a consequence relapse of disease occurs frequently. A thorough understanding of how LSCs self-renew and maintain their quiescent state is therefore essential in order to be able to develop targeting approaches that also show efficacy in those cell populations. We and others have shown that Polycomb group (PcG) proteins are important regulators of hematopoietic stem cell fate, both in health and in leukemia [1, 7–12]. PcG proteins are epigenetic regulators that are critically involved in controlling gene transcription by mediating post-translational modifications of histone proteins and chromatin remodeling [12–15]. Genome-wide analyses of Polycomb target genes revealed the occupancy of PcG proteins at promoters of genes regulating cell fate, highlighting their importance for proper lineage specification [16–18]. Yet, how PcG proteins are recruited, recognize their target genes and regulate gene expression still remains poorly understood. Understanding these processes is important since deregulation of PcG proteins frequently contributes to cancer and hematopoietic malignancies, like leukemia [19–21].

PcG proteins form multi-protein chromatin modifying complexes of which Polycomb Repressive Complex 1 (PRC1) and 2 (PRC2) are best characterized [22–25]. We recently identified an essential role for non-canonical PRC1.1 proteins in human leukemias [1]. PRC1.1 was first identified by the purification of the BCOR protein, which was found to interact with RING1A/B, RYBP, PCGF1, SKP1 and KDM2B [26, 27]. A potential oncogenic role of PRC1.1 is underlined by the fact that KDM2B is overexpressed in leukemias, breast and pancreatic cancers where it functions as an oncogene, conversely knockdown of KDM2B abrogated tumorigenicity [1, 28–31]. The exact function of individual subunits in the PRC1 complex is not fully understood, though it is suggested that they are involved in maintaining the integrity of the complex, in providing or controlling enzymatic activity or in targeting to chromatin [32–34]. For example, the H2AK119 E3 ligase activity is enhanced by the dimerization of RING1A/B with either PCGF2 or PCGF4 [35–37] or can be stimulated by the PCGF1-RYBP/YAF2 interaction in the case of non-canonical PRC1.1 [23, 34]. A shRNA approach for individual PRC1 subunits in hematopoietic stem cells revealed a lack of functional redundancy, suggesting unique functions of distinct PRC1 complexes [10] and indeed PRC1 complex composition changes upon lineage specification [38].

Protein ubiquitination is an important post-translational modification that controls the stability of almost all cellular proteins. Mono-ubiquitination impacts on the activity of proteins or can promote or prevent protein-protein interactions, while poly-ubiquitinated proteins are typically targeted to and degraded by the proteasome [39]. USP7 is a ubiquitin-specific protease that displays a wide range of activities, making it an attractive candidate target for cancer treatment. USP7 inhibition destabilizes MDM2 resulting in increased levels of TP53, and recently a number of USP7-specific inhibitors were generated that effectively targeted various human cancer cells presumably in an TP53-dependent manner [40–42]. However, TP53-independent roles exist as well [43, 44]. Here, using LC-MS/MS-based proteome studies we identify USP7 as an integral component of the KDM2B/PRC1.1 complex. USP7 inhibition results in PRC1.1 complex disassembly and reduced chromatin binding, with a concomitant reduction in gene expression of target loci. Our data show that USP7 is essential for leukemic cells and suggests that targeting of USP7 might provide an alternative therapeutic approach for leukemia, also for the most aggressive subtypes of AML which harbor mutations in TP53.

## MATERIALS AND METHODS

### GFP-mediated pull outs

Pull outs were performed on nuclear extracts from K562 cells stably expressing KDM2B-EGFP, PCGF1-EGFP and EGFP-RING1B. At least 80×10^6^ cells were collected for each pull out, nuclear extract preparation was done as described previously [10]. Pre-clearing of cell lysates was done by adding 50 µl pre-equilibrated binding control magnetic agarose beads (Chromotek) and incubated for 30 min at 4°C on a rotating platform. Then pre-cleared lysate was incubated with 70 µl pre-equilibrated GFP-Trap magnetic agarose beads (Chromotek) overnight at 4°C on a rotating platform. Beads were separated using a magnetic rack and six times washed in wash buffer (TBS, 0.3% IGEPAL CA-630, 1x CLAP, 0.1 mM PMSF). Bound fractions were eluted from the beads by boiling for 10 min in 2x Laemmli sample buffer.

### Chromatin immunoprecipitation

ChIP was essentially performed as described previously [45]. K562 cells stably expressing low levels of EGFP-fusion vectors encoding, PCGF1-EGFP, EGFP-RING1B, KDM2B-EGFP or non-transduced K562 cells and CB MLL-AF9 cells were treated with DMSO or P22077 for indicated timepoints and subsequently cross-linked. The following antibodies were used: anti-GFP (ab290, Abcam), anti-KDM2B (ab137547, Abcam), anti-H2AK119ub (D27C4, Cell Signaling Technology), anti-H3K4me3 (ab8580, Abcam), anti-H3K27ac (C15410196, Diagenode) and IgG (I8141, Sigma). ChIPs were analysed by qPCR as percentage of input, Supplementary Table 7 lists the qPCR primers used.

**Supplementary information is available online**

### Data sharing statement

For original data, please contact

## RESULTS

### Targeting of quiescent and cycling primary AML cell populations upon USP7 inhibition

To study the functional consequences of inhibition of the ubiquitin-specific protease 7 (USP7), we tested P22077 (1-(5-((2,4-difluorophenyl)thio)-4-nitrothiophen-2-yl)ethanone) on a panel of AML cell lines and primary patient samples. USP7 inhibition severely impaired cell growth in all tested AML cell lines (Figure 1A). Since P22077 has been documented to inhibit USP47 as well [46] we also tested the more specific USP7 inhibitor FT671 [40] which demonstrated similar reduced proliferation of MOLM-13, OCI-AML3, K562 and HL60 cells (Figure 1B). Cell lines that do not express functional TP53 (HL60 and K562, Supplementary Table 1) also showed sensitivity, indicating that at least in those cell lines the effect of P22077 and FT671 was independent of TP53. A strong dose-dependent reduction in cell viability and increase in apoptosis was observed as determined by Annexin V staining, while no significant changes in cell cycle distribution were noted (data not shown). Next, we assessed whether USP7 inhibition also affected the survival of primary AML cells grown in long-term stromal cocultures. The long-term proliferation of primary AML CD34^+^ cells was clearly reduced in the presence of P22077 (Figure 1C). Patient characteristics are shown in Supplementary Table 1, and again also a patient sample with a TP53(H179R) mutation showed strong sensitivity (Fig. 1c, AML 1). The more USP7-specific inhibitor FT671 was also tested on primary AML samples together with the somewhat less potent FT827 inhibitor [40], and again we observed strong reductions in proliferation, in particular in response to FT671 (Figure 1D).

**Figure 1:**
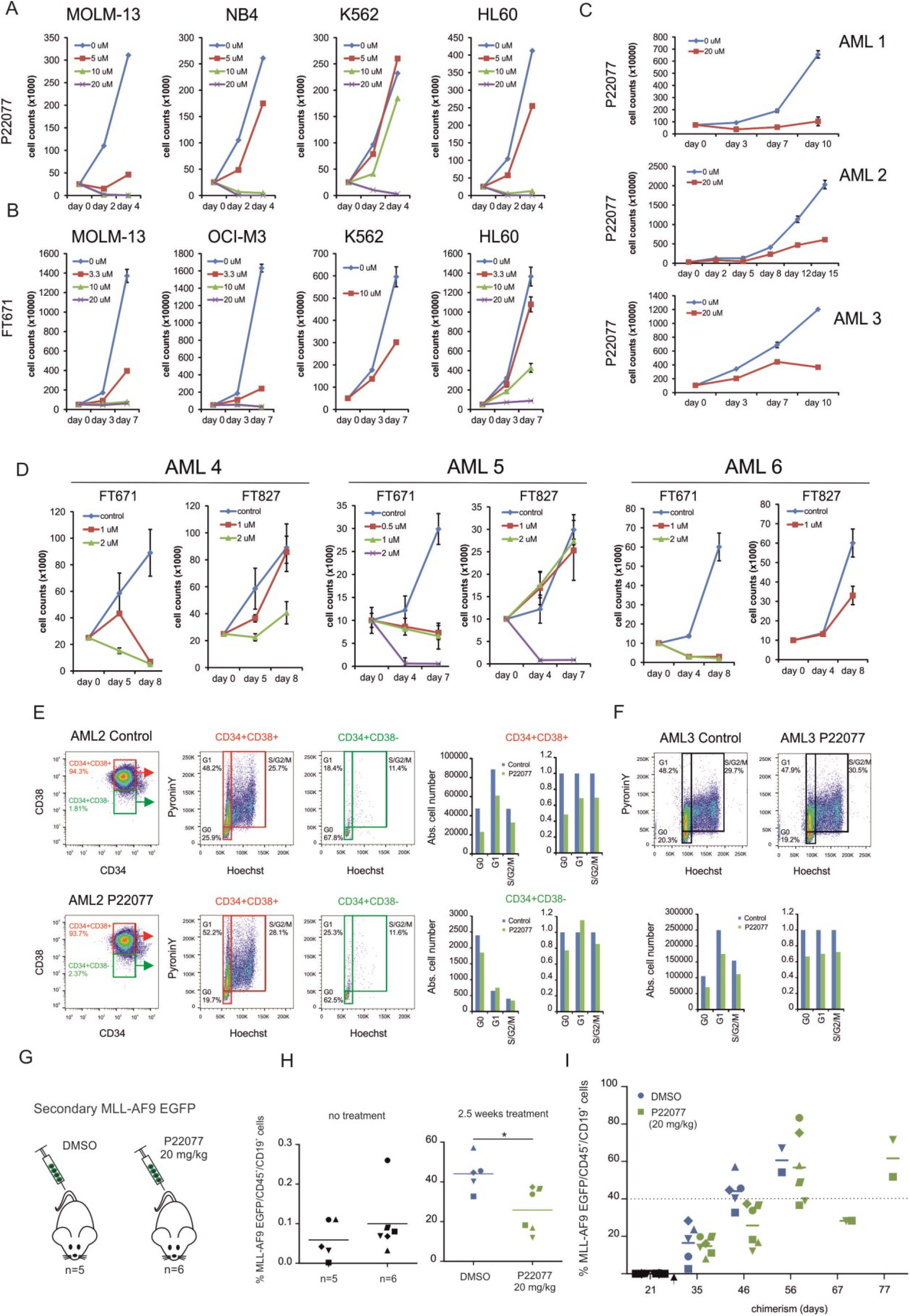
Sensitiviy of AML cells towards USP7 inhibition. (A) Cumulative cell growth of various AML cell lines in control (DMSO) or P22077 treated conditions. (B) Cumulative cell growth of various AML cell lines in control (DMSO) or FT671 conditions performed in triplicate. (C) Cumulative cell growth of primary AML patient cells (n=3) grown on MS5 co-culture treated with DMSO (control) or P22077 performed in duplicate. (D) Cumulative cell growth of primary AML patient cells (n=3) grown on MS5 co-culture treated with DMSO or FT671 performed in duplicate. (E-F) Analysis of both cycling and quiescent AML cells by Hoechst/PyroninY stainings within the CD34^+^/CD38^+^ and CD34^+^/CD38^−^ populations (E) or whole population (AML3) (F). Cells were treated with DMSO (control) or P22077 (20 µM) for 3 days while AMLs were grown on MS5 co-culture. Absolute cell numbers were calculated in G_0_, G_1_ and S/G_2_/M fractions. (G) Experimental setup of our human CB MLL-AF9 xenograft mouse model. Here 5 × 10^4^ MLL-AF9 EGFP cells from a primary leukemic mouse were IV injected into secondary recipients (n=11). (H) Peripheral blood analysis of MLL-AF9 EGFP/CD45^+^/CD19^+^ cells, three weeks after injection prior to treatment (left) and 2.5 weeks following treatment (right). Mice were treated daily with either DMSO as control (n=5) or 20 mg/kg P22077 (n=6). (I) Peripheral blood chimerism levels of control and P22077 (20 mg/kg) treated mice over the course of the experiment. Treatment was started at day 28 (4 weeks post-transplant) indicated with the arrow.

With current therapies for leukemia patients the majority of the blast population is readily eradicated, while the rare quiescent population of leukemic stem cells is more difficult to target. To examine whether USP7 inhibition does target the quiescent (G0) leukemic cell population we performed Hoechst/PyroninY stainings. AML 2 was characterized by a large CD34^+^/CD38^+^ population that is high in cycling activity, while the CD34^+^/CD38^−^ population was predominantly in G_0_ (Figure 1E). In both populations, the G_0_ fraction was equally or even more efficiently targeted by P22077 compared to the cycling fraction, indicating that cycling cells are not necessarily more sensitive than quiescent cells. AML3 was low in CD34^+^ and therefore we gated on the whole blast population which also showed that both quiescent and cycling cells were sensitive for P22077 (Figure 1F).

Next, we evaluated the effect of USP7 inhibition in our human CB MLL-AF9 xenograft mouse model (Figure 1G) [47, 48]. Importantly, these lymphoid MLL-AF9 cells when grown in co-culture on MS5 stroma *in vitro* were sensitive towards USP7 inhibition (data not shown). 5 × 10^4^ MLL-AF9 leukemic cells, from a primary leukemic mouse, were intravenously (IV) injected into secondary recipients (n=11). Three weeks after injection 10/11 mice showed engraftment of MLL-AF9 EGFP^+^/CD45^+^/CD19^+^ cells in peripheral blood and subsequently mice were divided into two groups that were treated either with DMSO (n=5) or 20 mg/kg P22077 (n=6) via intraperitoneal (IP) injections. Mice were treated daily starting four weeks post-transplant and peripheral blood chimerism levels were monitored by regular blood sample analysis and mice were sacrificed when chimerism levels in the blood exceeded 40%. Two-and-a-half weeks after the initiation of treatment chimerism levels were significantly lower in P22077-treated mice compared to DMSO (Figure 1H). The chimerism levels for DMSO treated mice rapidly increased to 44% (average) within 6 weeks after injection, whereas the chimerism levels of P22077 treated mice were around 25% on average. Notably, 3/6 mice gave a better response to USP7 inhibition with lower chimerism levels of 18.3, 16.8 and 12% respectively at day 46 (Figure 6I). Within 60 days all control mice exceeded 40% chimerism in the blood, indicative for a full blown leukemia and were sacrificed (Figure 6I). Bone marrow, spleen and liver analyses showed high levels (>90%) of chimerism (data not shown). Leukemia development was significantly delayed in USP7 inhibitor treated mice, and in particular in two mice a clear response to USP7 inhibition was observed and chimerism levels remained relatively stable between day 56-67 post-transplant, although ultimately those mice also did develop MLL-AF9-induced leukemia after day 77 (Figure 1I).

### Identification of the deubiquitinase USP7 as a subunit of PRC1.1

One of the pathways downstream of USP7 via which its inhibition might contribute to reduced cell survival is the TP53 pathway [49]. Yet, also in the absence of functional TP53, both in leukemic cell lines as well as in primary patient samples, we noted strong sensitivity towards USP7 inhibition. We recently identified KDM2B as a critically important factor for the survival of human leukemic stem cells [1] and in our initial interactome screens we had already identified USP7 as a potential interaction partner within the non-canonical PRC1.1 complex. Therefore, we set out to further study the KDM2B interactome in detail in leukemic cells, specifically focussing on a potential role for USP7. Human K562 cells were transduced with KDM2B-EGFP followed by anti-EGFP pull outs and LC-MS/MS analysis to identify interacting proteins. Thus, 406 KDM2B-interacting proteins were identified involved in cellular processes like DNA/chromatin binding, protein binding, RNA binding and RNA polymerase activity (Figure 2A and Supplementary Table 2). Gene Ontology analyses revealed that this list contained proteins that associated with rRNA processing, mRNA splicing, translation, mRNA processing, positive regulation of gene expression and DNA damage response (Figure 2B). The most abundant KDM2B interaction partners were non-canonical PRC1.1 proteins (Figure 2C and 2D), including USP7. Next, we questioned whether inhibition of the deubiquitinase activity of USP7 would impact on PRC1.1 function, which would provide further insight into possible mechanisms via which USP7 inhibitors would contribute to targeting of leukemic (stem) cell populations. To further validate the interaction of USP7 with PRC1.1 proteins, besides KDM2B, we also performed EGFP pull outs on nuclear extracts of stable K562 cell lines expressing PCGF1-EGFP and EGFP-RING1B followed by LC-MS/MS analysis (Supplementary Table 3). Figure 2D shows the spectral counts (MS/MS counts) corrected for expected peptides based on *in silico* protein digests. Clearly, USP7 was found to interact with KDM2B, PCGF1 and RING1B and all other PRC1.1 proteins were co-purified, indicating that USP7 is a bona fide subunit of PRC1.1. Importantly, USP47 was not found as interaction partner with any of the Polycomb proteins (Supplementary Table 2-3). In addition, Western blots of independent pull outs on canonical PRC1 (PCGF2, PCGF4, CBX2) and non-canonical PRC1.1 (PCGF1, RING1B) were performed using antibodies against USP7 and RING1B (Figure 2E). As expected, RING1B precipitated with both PRC1 and PRC1.1 proteins. The interaction of USP7 with RING1B and PCGF1 was confirmed by Western blot while less interaction was observed with the canonical PRC1 proteins PCGF2, PCGF4 or CBX2 (Figure 2E and Supplementary Table 4).

**Figure 2.**
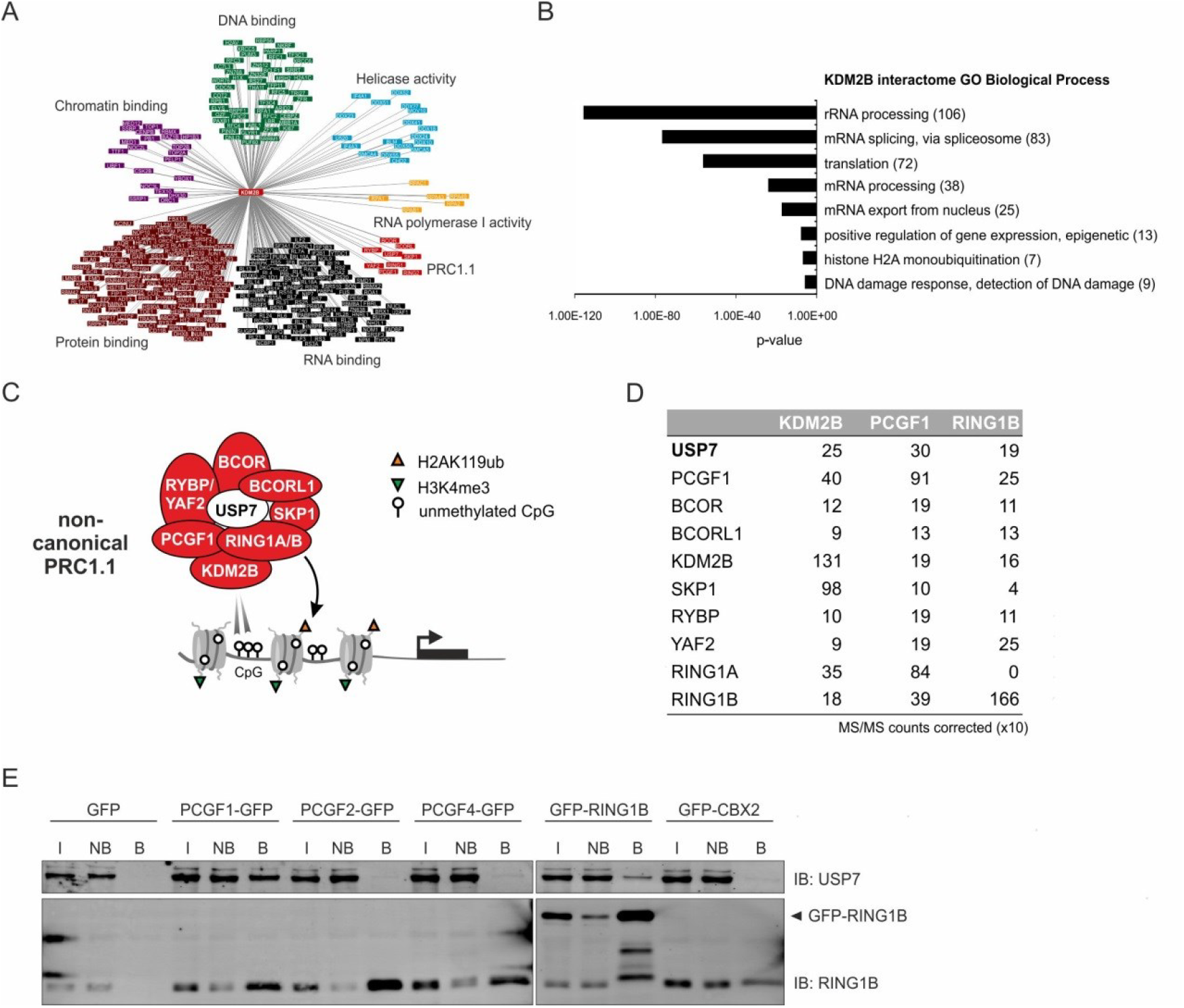
The deubiquitinase USP7 interacts with non-canonical PRC1.1 proteins. (A) Visualization of KDM2B interactome using Cytoscape, categorized by Molecular Function as annotated by DAVID. (B) Gene Ontology analysis of KDM2B interactome shown as Biological Process. (C) Schematic representation of the non-canonical PRC1.1 complex, including USP7. PRC1.1 preferentially binds to non-methylated CpG islands via the ZF-CxxC domain of KDM2B. The ubiquitination of histone H2A on lysine 119 is mediated by RING1A/B. PRC1.1 can be recruited to active genes in leukemic cells, indicated by H3K4me3. (D) Identification of USP7 and other PRC1.1 proteins by LC-MS/MS in KDM2B-EGFP, PCGF1-EGFP and EGFP-RING1B pull outs from nuclear extracts in K562 cells. The numbers indicate MS/MS counts for each interacting protein corrected for expected peptides based on *in silico* digests (x10). (E) Validation of USP7 and RING1B interactions as analysed by Western blot in canonical PRC1 (PCGF2/PCGF4/CBX2/RING1B) and non-canonical PRC1.1 (PCGF1/RING1B) pull outs. Input (I), non-bound (NB) and bound (B) fractions are shown.

### USP7 inhibition results in disassembly of the PRC1.1 complex

Deubiquitinating enzymes (DUBs) exert a variety of important cellular functions, including the control over protein stabilization or degradation, protein localization, protein activity or by modulating protein-protein interactions [50]. We therefore questioned whether inhibition of the deubiquitinase USP7 might affect PRC1.1 stability and function. In general, DUB inhibitors increase overall protein polyubiquitination of many target proteins [51]. Similarly, we observed accumulation of polyubiquitinated proteins in our EGFP-RING1B and PCGF1-EGFP K562 cells treated for 24h with P22077 (Figure 3A). Next, we investigated the effect of USP7 inhibition on the stability of PRC1.1. EGFP pull outs were performed on nuclear extracts from K562 PCGF1-EGFP and EGFP-RING1B cells treated with DMSO or P22077 followed by LC/MS-MS analysis (Figure 3B and Supplementary Table 5). Volcano plots were generated using label-free quantification (LFQ) intensities of potential interactors of EGFP-RING1B (left) and PCGF1-EGFP (right) plotted as fold change difference (control/USP7i) against significance (t-test p-value). Interactions with several PRC1.1 proteins, highlighted in orange, were significantly reduced in both EGFP-RING1B and PCGF1-EGFP pull outs as a consequence of USP7 inhibition (Figure 3B). The ubiquitin protein, UBB, was enriched in both PCGF1-EGFP and EGFP-RING1B pull outs upon P22077 treatment, suggesting that these proteins or other Polycomb interaction proteins might be more ubiquitinated upon USP7 inhibition. In addition we analysed the intensity-based absolute quantification (iBAQ) values, as a measure for protein abundance, relative to either RING1B or PCGF1 and normalized to control pull outs (Figure 3C). Similarly, these data demonstrated a clear reduced interaction of PRC1.1 proteins with RING1B and PCGF1 after P22077 treatment. Independent EGFP pull outs performed on PCGF1-EGFP and EGFP-RING1B cell lines further confirmed that USP7 inhibition indeed resulted in reduced interaction of PCGF1-EGFP with endogenous RING1B and EGFP-RING1B with endogenous PCGF1 (Figure 3D). Importantly, input samples did not reveal reduced expression of EGFP-RING1B,PCGF1-EGFP or KDM2B-EGFP, which was further validated by FACS analysis (Figure 3E). Taken together these data indicate that USP7 is essential for PRC1.1 complex integrity.

**Figure 3.**
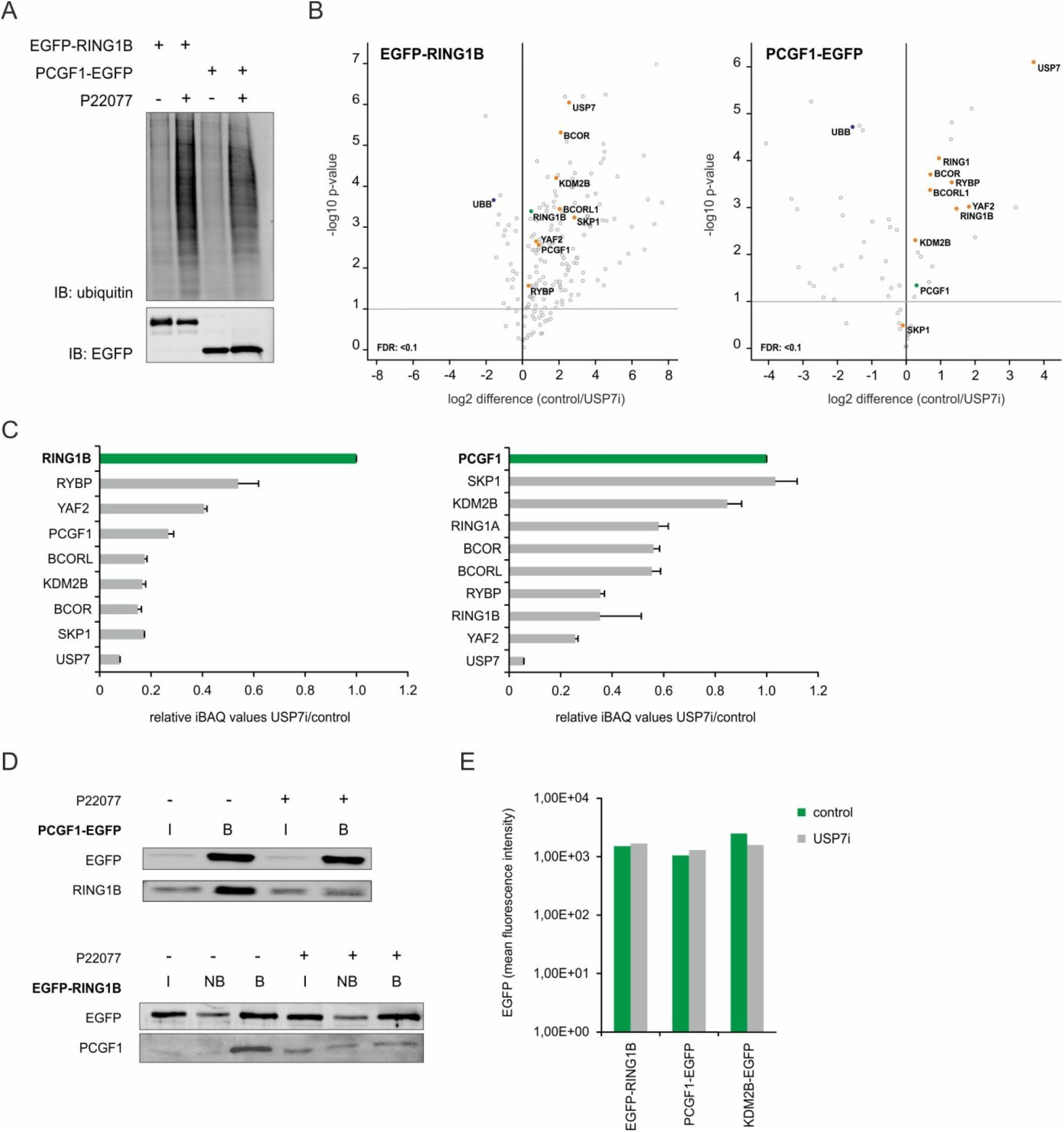
USP7 deubiquitinase activity is essential for PRC1.1 integrity. (A) Purification of his-tagged ubiquitinated proteins under denaturing conditions in EGFP-RING1B and PCGF1-EGFP cells treated with DMSO (-) or P22077 (+) for 24h followed by Western blot analysis. (B) Label-free quantification (LFQ) intensities of EGFP pull outs on EGFP-RING1B (left) and PCGF1-EGFP (right) interactions, performed in technical replicates in control (DMSO, 0.1%) and USP7i treated cells are illustrated as Volcano plot. The fold change difference (log2) of the EGFP pull out in control versus USP7i is plotted against the –log10 t-test p-value (y-axis). The bait protein (RING1B/PCGF1) is highlighted in green, PRC1.1 proteins in orange and UBB protein in purple. (C) Intensity based absolute quantification (iBAQ) values for several identified PRC1.1 proteins are shown in USP7i/control EGFP pull outs relative to RING1B (left) and PCGF1 (right). Data are shown as mean ± SD (n=2 or 3). (D) Western blot of EGFP pull outs on PCGF1-EGFP and EGFP-RING1B in the absence (-) or presence (+) of P22077 for 72h probed with antibodies for EGFP, RING1B and PCGF1. Input (I), non-bound (NB) and bound (B) fractions are shown. (E) Mean fluorescent intensity (MFI) analysis of EGFP-RING1B, PCGF1-EGFP AND KDM2B-EGFP K562 cells in control or USP7i treated cells at 72h.

### PRC1.1 chromatin binding relies on functional USP7 deubiquitinase activity

To determine the functional consequences of disassembly of the PRC1.1 complex upon USP7 inhibition, we performed several ChIPs to investigate whether PRC1.1 chromatin targeting at target loci would be affected by USP7 inhibition. Previously, we identified PRC1 and PRC1.1 target loci by ChIP-seq for PCGF1/2/4, CBX2, RING1A/1B, KDM2B, H2AK119ub and H3K27me3 in the leukemic cell line K562 [1]. Here, ChIP-qPCRs were performed for KDM2B, PCGF1 and RING1B, the core PRC1.1 subunits, on several loci in the absence or presence of P22077. Inhibition of USP7 resulted in a complete loss of KDM2B binding, with concomitant strong reductions in PCGF1 and RING1B chromatin binding (Figure 4A). Since RING1B mediates H2AK119 ubiquitination we then analysed the levels of H2AK119ub upon USP7 inhibition. Similarly, H2AK119ub was lost from PRC1.1 target loci (Figure 4B). These data indicate that USP7 inhibition severely impacts on PRC1.1 chromatin binding and as a consequence on H2AK119ub levels. Since non-canonical PRC1.1 can be recruited to active loci in leukemic cells, we also investigated the levels of H3K4me3, but no changes were observed on this histone mark upon USP7 inhibition, highlighting that not all posttranslational histone modifications are affected as a consequence of treatment (Figure 4C). To further investigate the kinetics of loss of H2AK119ub cells were cross-linked at 4h, 8h and 16h followed by a ChIP for H2AK119ub, KDM2B and RING1B (Figure 4D). The time-dependent loss of H2AK119ub coincided with reduced KDM2B and RING1B binding at the same target loci, suggesting that loss of *de novo* ubiquitination underlies these observations. Again, H3K4me3 levels remained unaffected, while H3K27ac levels were reduced upon USP7 inhibition indicative for reduced transcriptional activity at these loci (Figure 4E). Finally, to exclude a potential role of USP47 we also validated our findings using the more specific USP7 inhibitor FT671, and these data also clearly demonstrated a loss of KDM2B binding and reduced H2AK119ub levels at PRC1.1 loci upon inhibiting USP7 function (Figure 4F).

**Figure 4.**
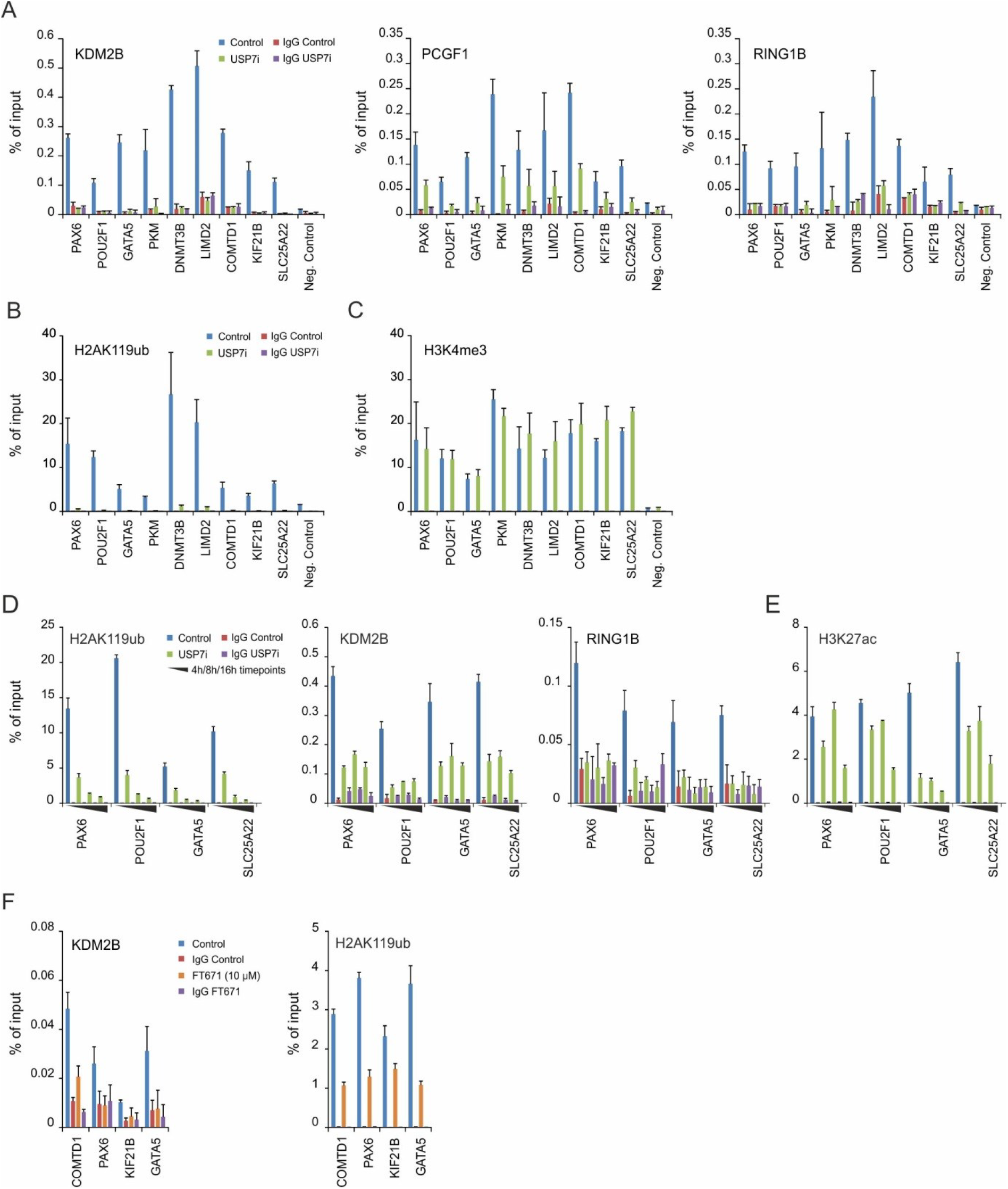
Loss of PRC1.1 occupancy and H2AK119ub at target loci upon USP7 inhibition. (A) ChIP-qPCRs on control or 72h P22077 (USP7i) treated K562 cells for endogenous KDM2B, PCGF1-EGFP, EGFP-RING1B (B) H2AK119ub and (C) H3K4me3 on several PRC1.1 loci previously identified by ChIP-seq. Error bars represent SD of technical qPCR replicates. (D) ChIP-qPCRs on control or USP7i treated K562 cells for 4h/8h/16h 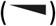 for H2AK119ub, KDM2B-EGFP, EGFP-RING1B and (E) H3K27ac on four PRC1.1 loci. Error bars represent SD of technical qPCR replicates. (F) ChIP-qPCRs on control or 24h FT671 (10 µM) treated K562 cells for endogenous KDM2B and H2AK119ub. Error bars represent SD of technical qPCR replicates.

### USP7 inhibition leads to downregulation of several PRC1.1 active target genes

Given that USP7 inhibition severely impaired PRC1.1 occupancy to several target loci we investigated whether this would also affect the expression of PRC1.1 target genes in leukemic cells. RNA-seq was performed to compare gene expression in DMSO or USP7 inhibitor treated K562 cells (P22077, 30 µM) for 4h, 8h, 16h and 24h. Since PRC1.1 can be associated with active genes in leukemic cells we focused on the genes that were downregulated upon USP7 inhibition. In total 297 genes were downregulated more than 2 fold after 24h of USP7 inhibition (Supplementary Table 6). Since USP7 has also PRC1.1 independent functions, for instance by controlling MDM2/TP53 [52], we then analysed whether downregulated genes overlapped with previously identified PRC1.1 peaks by ChIP-seq (Figure 5A and Supplementary Table 6)[1]. Twenty-four percent of these genes overlapped with PRC1.1 targets. Gene Ontology (GO) analysis revealed that this set was enriched for GO terms like transcription, regulation of gene expression and chromatin modification, while PRC1.1 independent downregulated genes were enriched for GO terms like mRNA splicing, protein folding and protein polyubiquitination (Fig. 5a). ChIP-seq profiles of TOP2B, SIN3A, CHD1 and MYC are shown in Fig. 5b as representative examples for PRC1.1 target genes. The tracks show clear binding of PCGF1, RING1A/1B and KDM2B and little occupancy of canonical PRC1 proteins (PCGF2, PCGF4, CBX2). These loci were also enriched for H2AK119ub and active chromatin marks H3K4me3, H3K36me3, RNAPII and H3K27ac but devoid of the repressive mark H3K27me3. Loss of PRC1.1 binding was confirmed by ChIP-qPCRs for KDM2B, PCGF1 and RING1B on these 4 loci (Figure 5C). Again, a concomitant loss of H2AK119ub and H3K27ac marks was seen, without any alterations in H3K4me3 (Figure 5C). Subsequently, the observed downregulation based on RNA-seq data was validated by independent quantitative RT-PCRs (Figure 5D). Loss of PRC1.1 binding correlated with loss of gene expression. Taken together, these data suggest that the presence of PRC1.1 is required to maintain the transcriptional activity of several target genes.

**Figure 5.**
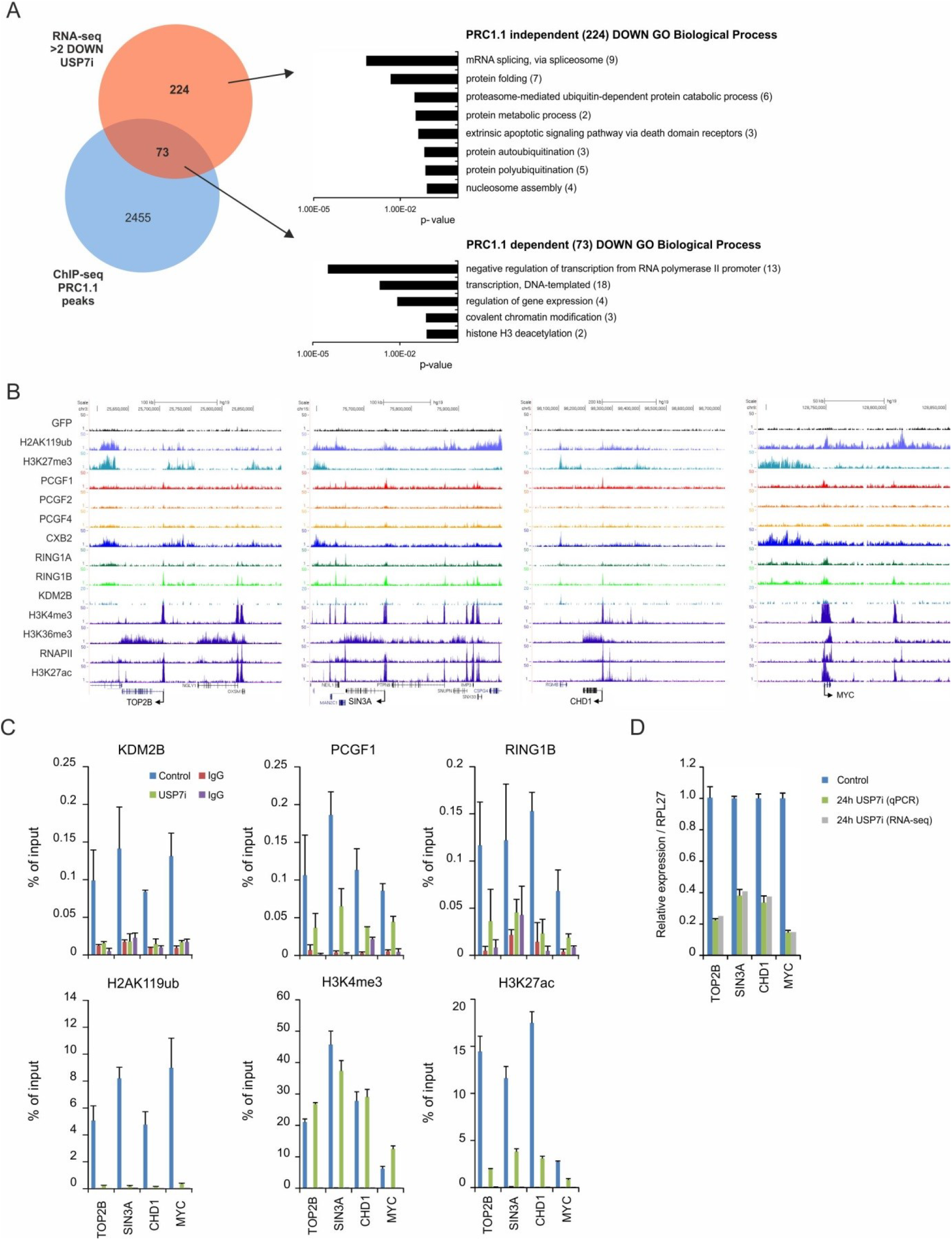
USP7 inhibition leads to downregulation of a subset of PRC1.1 active target genes. (A) Venn diagram showing overlap of genes downregulated by more than two fold after 24h of USP7 inhibition with previously identified PRC1.1 target genes. Gene Ontology analysis of downregulated genes identified as PRC1.1 targets (73) or Polycomb independent targets (224) shown as Biological Process. (B) ChIP-seq profiles of TOP2B, SIN3A, CHD1 and MYC as representative examples for PRC1.1 target genes (73). Our ChIP-seq tracks are shown for GFP (control), H2AK119ub, H3K27me3, PCGF1, PCGF2, PCGF4, CBX2, RING1A, RING1B and KDM2B. ChIP-seq tracks for H3K4me3, H3K36me3, RNAPII and H3K27ac were downloaded from ENCODE/Broad. (C) ChIP-qPCRs in control or 72h USP7i for KDM2B, PCGF1-EGFP, EGFP-RING1B and H2AK119ub or 16h USP7i for H3K4me3 and H3K27ac. Error bars represent SD of technical qPCR replicates. (D) Validation of relative gene expression levels of PRC1.1 target genes by qRT-PCR in control and P22077 treated cells (24h). Data are represented as mean ± SD.

## DISCUSSION

In this study, we reveal that the deubiquitinase USP7 is essential for leukemic cells and suggests that targeting of USP7 might provide an alternative therapeutic approach for leukemia, also for the most aggressive subtypes of AML which harbor mutations in TP53. Mechanistically, using these TP53null AML cells as starting point, we uncover an important role for USP7 in controlling the stability and function of non-canonical PRC1.1 in leukemia. Our interactome proteomics studies identify USP7 as an integral component of PRC1.1, and inhibiting USP7 deubiquitinase activity resulted in disassembly of the PRC1.1 complex. Consequently, recruitment of the PRC1.1 complex to target loci was lost coinciding with the loss of its H2AK119ub catalytic activity. This resulted in repression of a subset of transcriptionally active PRC1.1 target genes and corresponded with a reduction in H3K27ac, highlighting an essential function for PRC1.1 in maintaining gene transcription.

Our data indicates that USP7 stably interacts with PRC1.1 proteins, KDM2B, PCGF1 and RING1B (Fig. 1). Sánchez and colleagues first identified the ubiquitin protease USP7 as an interactor of RING1B [27]. Then, USP7 was also identified in proteomics analysis of PCGF1, RYBP, YAF2 and RING1A/B pullouts [1, 23, 53]. Moreover, using quantitative proteomics and USP7 as bait specific interactions were identified in HeLa cells with PCGF1, BCOR, RING1A/B [53]. Previously, USP7 has been shown to be associated with the canonical PRC1 protein BMI1 (PCGF4) and potentially also with MEL18 (PCGF2), although Lecona and colleagues suggested that USP7 interacts directly with SCML2 and thereby bridges the interaction with PRC1.4 [54, 55]. However, similar to Gao and colleagues [23], our PCGF2, PCGF4 and CBX2 proteome analyses showed little or no interaction with USP7 [1]. We did notice that several canonical RING1B-Polycomb interactions were also reduced upon USP7 inhibition, including interactions with CBX2/4/8, SCML2 and SCMH1, though the interaction with PCGF2 and PCGF4 was not affected. This might also suggest that USP7 impacts on PRC1 protein stability, although further studies are required to clarify these issues. In line with that notion, a recent paper demonstrated that another deubiquitinase, USP26, controls the stability of CBX4 and CBX6 and thereby affects the complex composition of PRC1 during ESC reprogramming [56]. Maertens and colleagues demonstrated that also USP11 interacts with PRC1 proteins and affects the stability of BMI1 [55]. Thus, DUBs clearly play an important role in controlling Polycomb stability and complex composition.

Given that USP7 inhibition affects the complex composition of PRC1.1, its activity might control ubiquitin levels of Polycomb proteins, important for protein-protein interactions within PRC1.1. Upon USP7 inhibition an overall increase in protein ubiquitination was observed (Figure 2). Importantly, USP7 inhibition did not lead to degradation of PCGF1 or RING1B, allowing us to conclude that USP7 controls protein-protein interactions rather than proteasomal degradation. Overexpression of USP7 has been shown to deubiquitinate RING1B thereby stabilizing RING1B and preventing it from proteasomal degradation [57]. Furthermore, RING1B self-ubiquitination is required for its ligase activity on histone H2A [58]. This is supported by the observation that H2AK119ub is lost upon USP7 inhibition, most likely as a consequence of loss of *de novo* ubiquitination mediated via RING1B. Similarly, knockout of USP7 reduced H2AK119ub levels coinciding with reduced levels of RING1B [54]. Interestingly, UbE2E1 was shown to be a essential for Polycomb-mediated H2AK119ub and regulated by USP7 [59, 60]. Other studies also show that USP7 is not likely a DUB for H2AK119 [55, 61, 62]. Thus, regulating ubiquitin levels within the PRC1.1 complex itself is critically important and controlled by USP7 deubiquitinase activity. Inhibiting the catalytic core by removal of RING1A/B in ESCs does not lead to loss of KDM2B binding [63]. However, in leukemic cells we find that USP7 inhibition not only leads to reduced H2AK119ub but also induces loss of PRC1.1/KDM2B binding as a result of complex disassembly, suggesting that KDM2B-PCGF1-BCOR/L1/SKP1 interactions might be needed for targeting. This is in agreement with data that suggest that dimerization of PCGF1 and BCOR(L1) is required for binding to KDM2B and recruiting PRC1.1 to the chromatin [64].

With the efficient loss of PRC1.1 binding and consequent loss of H2AK119ub from target loci upon USP7 inhibition we were able to study PRC1.1 function in relation to gene regulation in more detail. Previously we identified genes that are targeted by PRC1.1 independent of H3K27me3 and have a transcriptionally active profile suggested by H3K4me3, RNAPII and H3K27ac occupancy close to the transcription start site. Loss of PRC1.1 binding resulted in reduced gene expression coinciding with reduced levels of H3K27ac on several known PRC1.1 targets. We therefore hypothesize that PRC1.1 creates a transcriptionally permissive and open chromatin state which enables transcription factors to bind and initiate gene expression. Addressing a possible cross-talk between PRC1.1 proteins, CBP/p300-linked H3K27ac, recruitment of transcription factors and accessibility of chromatin is definitely a focus for future work. Since H3K4me3 levels remained unaffected following USP7 inhibition, this suggest that the H3K4 methyltransferase complex that is likely targeted to CpGs via CFP1 protein is not targeted in a PRC1.1-dependent manner [65].

Where we previously highlighted the importance of PRC1.1 for the survival of leukemic cells using genetic studies, small molecule inhibitors would be easier to implement in a therapeutic clinical setting [1]. Inhibition of USP7 provided an efficient means to target PRC1.1. Of course it is evident that USP7 can function in several pathways, often through regulating protein stability of tumor suppressors or epigenetic regulators [66–68] and it is particularly the TP53 pathway that is strongly controlled by USP7 [49, 69–71]. Various USP7 inhibitors have been developed, and most recently selective USP7 inhibitors were generated that destabilize USP7 substrates including MDM2 and thereby increase TP53-mediated apoptosis of cancer cells [40, 72]. While we cannot exclude the possibility that some of these pathways were also affected in some of our studies, we have analysed the TP53 status in our models and primary AML patient samples and see sensitivity even in the absence of a normal TP53 response. Furthermore, while USP7 knockout mice are embryonically lethal, deletion of p53 was not able to rescue this phenotype, further highlighting that p53-independent pathways downstream of USP7 exist as well [73, 74].

In conclusion, our data reveal an important role for USP7 deubiquitinase activity in the integrity of the PRC1.1 complex. We provide insight into the recruitment of PRC1.1 to target loci and function in gene regulation and show that PRC1.1 is a potential interesting therapeutic target in leukemia.

## Acknowledgements

The authors thank Dr. S. Bergink and E. de Mattos for providing pCl His-Ubi plasmid and advice with ubiquitin experiments. We thank M. Elliot for the generation of KDM2B-EGFP construct. We also thank Bart-Jan Wierenga for generation of the pRRL.SFFV.EGFP.miR-E vector. The authors thank FORMA therapeutics (Watertown, MA, USA) for providing the FT671 and FT827 compounds. We would like to thank Roelof-Jan van der Lei, Theo Bijma, Henk Moes en Geert Mesander for help with cell sorting. We acknowledge Prof. Robert E Campbell (Department of Chemistry, University of Alberta, Edmonton, Alberta, Canada) for providing the mBlueberry2 Fluorescent Protein. This work is supported by the European Research Council (ERC-2011-StG 281474-huLSCtargeting) and the Dutch Cancer Foundation (RUG 2014-6832).

## Author contributions

HM designed and performed experiments, analyzed data, and wrote the manuscript; JJ, ARL, SMH, MPV, CG, AZBV, NM, VB performed experiments, analyzed data, and reviewed the manuscript; GH and EV provided patient samples, analyzed data and reviewed tha manuscript; JJS designed the study, analyzed data and wrote the manuscript.

