## Supplementary files and figures for "USP7 as part of non-canonical PRC1.1 is a druggable target in leukemia"

**Running title: USP7 maintains integrity and function of PRC1.1**

Henny Maat1, Jennifer Jaques1, Aida Rodríguez López1, Shanna M. Hogeling1, Marcel P. de Vries2,3, Chantal Gravesteijn1, Annet Z. Brouwers-Vos1, Nisha van der Meer1, Gerwin Huls1, Edo Vellenga1, Vincent van den Boom1 and Jan Jacob Schuringa1,*.

1Department of Experimental Hematology, Cancer Research Center Groningen, University Medical Center Groningen, University of Groningen, Hanzeplein 1, 9713 GZ Groningen, The Netherlands;

2Department of Pharmacy, Interfaculty Mass Spectrometry Center, University of Groningen, A. Deusinglaan 1, 9713 AV Groningen, The Netherlands;

3Department of Pediatrics, Center for Liver, Digestive, and Metabolic Diseases, University Medical Center Groningen, University of Groningen, Hanzeplein 1, 9713 GZ Groningen, The Netherlands

**Supplementary Methods**

**Cell culture**

The AML cell lines K562, HL60 (ATCC: CCL-243,CCL-240), MOLM13, NB4 and OCI-AML3 (DSMZ: ACC-554, ACC-207, ACC-582) were cultured in RPMI 1640 (BioWhittaker, Lonza, Verviers, Belgium) supplemented with 10% fetal bovine serum (FCS, HyClone Laboratories, Logan, Utah, US) and 1% penicillin/streptomycin (p/s, PAA Laboratories). MS5 murine stromal cells (DSMZ: ACC-441) were cultured in alpha-MEM with 200 mM glutamine (BioWhittaker) supplemented with 10% FCS and 1% p/s. Primary AMLs were cultured in Gartner's medium as described before [44]. CB MLL-AF9 liquid and MS5 co-cultures under myeloid or lymphoid permissive conditions were performed as described previously (1, 2). All cultures were kept at 37°C and 5% $CO_2$. For USP7 inhibition experiments, P22077 (1-(5-((2,4-difluorophenyl)thio)-4-nitrothiophen-2-yl)ethanone) was purchased from Merck Millipore (662142) (Billerica, MA, USA). FT671 and FT827 were generously provided by FORMA Therapeutics (Watertown, MA, USA (3).

**In gel trypsin digestion**

Bound fractions of KDM2B-EGFP, PCGF1-EGFP (control/USP7i) and GFP-RING1B (control/USP7i) were loaded on a 4-12% pre-cast NuPAGE gel (Invitrogen) and were run briefly. Gels were stained with Coomassie dye R-250 (Thermo Scientific) and subsequently destained with ultrapure water overnight. Gel lanes were cut into one slice, further cut into small pieces and completely destained using 70% 50 mM $NH_4HCO_3$ and 30% acetonitrile (ACN). Reduction and alkylation of cysteines was performed by adding 10 mM DTT dissolved in 50 mM $NH_4HCO_3$ and incubated at 55°C for 30 min. Next, 55 mM iodoacetamide in 50 mM $NH_4HCO_3$ was added and incubated for 30 min, in dark, at room temperature. Remaining fluid was removed and 50 mM $NH_4HCO_3$ was added with 10 min. shaking. Then 100% ACN was added incubated for 30 min. while shaking. Fluid was removed and gel pieces were dried for 15 min. at 55°C. Proteins were digested with adding 10 ng/µl sequencing-grade modified trypsin (Promega) in 50 mM $NH_4HCO_3$ to the gel pieces and incubated overnight at 37°C. Next day, peptides were extracted using 5% formic acid followed by second elution with 5% formic acid in 75% acetonitrile. Samples were dried in a SpeedVac centrifuge and dissolved in 5% formic acid.

**LC-MS/MS analysis**

Online chromatography of the extracted tryptic peptides was performed with the Ultimate 3000 nano-HPLC system (Thermo Fisher Scientific) coupled online to a Q-Exactive-Plus mass spectrometer with a NanoFlex source (Thermo Fisher Scientific) equipped with a stainless steel emitter. Tryptic digests were loaded onto a 5 mm × 300 µm i.d. trapping micro column packed with PepMAP100 5 µm particles (Dionex) in 0.1% FA at the flow rate of 20 µL/min. After loading and washing for 3 minutes, peptides were forward-flush eluted onto a 50 cm × 75 µm i.d. nanocolumn, packed with Acclaim C18 PepMAP100 2 µm particles (Dionex). The following mobile phase gradient was delivered at the flow rate of 300 nL/min: 2–50% of solvent B in 90 min; 50–80% B in 1 min; 80% B during 9 min, and back to 2 % B in 1 min and held at 3% A for 19 minutes. Solvent A was 100:0 $H_2O$/acetonitrile (v/v) with 0.1% formic acid and solvent B was 0:100 $H_2O$/acetonitrile (v/v) with 0.1% formic acid. MS data were acquired using a data-dependent top-10 method dynamically choosing the most abundant not-yet-sequenced precursor ions from the survey scans (300–1650 Th) with a dynamic exclusion of 20 seconds . Sequencing was performed via higher energy collisional dissociation fragmentation with a target value of 2e5 ions determined with predictive automatic gain control. Isolation of precursors was performed with a window of 1.6. Survey scans were acquired at a resolution of 70,000 at m/z 200. Resolution for HCD spectra was set to 17,500 at m/z 200 with a maximum ion injection time of 110 ms. The normalized collision energy was set at 28. Furthermore, the S-lens RF level was set at 60 and the capillary temperature was set at 250degr. C. Precursor ions with single, unassigned, or six and higher charge states were excluded from fragmentation selection.

**Data analysis**

Raw mass spectrometry data were analysed using MaxQuant version 1.5.2.8 (4), using default settings and LFQ/ iBAQ enabled, searched against the Human Uniprot/Swissprot database (downloaded June 26, 2016, 20197 entries). Further data processing was performed using Perseus software, version 1.5.6.0 (5). Network visualization of KDM2B interactome was performed using Cytoscape software, version 3.5.1. For Gene Ontology (GO) analysis we used DAVID Bioinformatics Resources (http://david.abcc.ncifcrf.gov/home.jsp). ChIP-seq tracks were visualized and analysed using UCSC genome browser (http://genome.ucsc.edu). Accession number for the Polycomb ChIP-seq data previously reported and used for analysis in this paper is GEO: GSE54580 (2). Previously published ChIP-seq used for analysis include H3K4me3, H3K36me3, H3K27ac and RNAPII/Pol2(b) from ENCODE/Broad Institute (GSE29611).

**Patient samples**

AML blasts from peripheral blood or bone marrow from untreated patients were studied after informed consent and the protocol was approved by the Medical Ethical Committee, in accordance with the Declaration of Helsinki. Mononuclear cells were isolated by density gradient centrifugation and CD34+ cells were selected automatically by using autoMACS (Miltenyi Biotec).

**Generation of lentiviral vectors and transductions**

Lentiviral pRRL SFFV PCGF1-EGFP and EGFP-RING1B vectors were generated as described previously(2). pRRL SFFV KDM2B EGFP was generated as follows. KDM2B was PCR amplified from cDNA in two parts (from the ATG to the RsrII site [fragment 1] and from the RsrII site to end of KDM2B [excluding the stop codon, fragment 2]). Both fragments were independently subcloned into pJet1.2 resulting in the pJet1.2 KDM2B[1] and pJet1.2 KDM2B[2] plasmids and that were subsequenty verified by sequencing. Next, KDM2B fragment 2 was isolated from pJet1.2 KDM2B[2] using RsrII and XbaI digestion and ligated into pJet1.2 KDM2B[1] that was also digested with RsrII and

XbaI, resulting in a pJet1.2 plasmid with the full length KDM2B ORF but excluding the stop codon. Finally, the KDM2B ORF was subcloned into the pRRL SFFV GFP vector using AgeI digestion, resulting in the pRRL SFFV KDM2B-GFP construct. A lentiviral pRRL SFFV-His-Ubiquitin-mBlueberry2 vector was generated by PCR amplification of His-Ubiquitin from pCl His-Ubi (Addgene) plasmid using primers including BamHI sites. The PCR product was first ligated into pJet1.2 using Blunt-End cloning protocol. Subsequently His-Ubiquitin was isolated from pJet1.2 using BamHI digestion and ligated into pRRL SFFV-IRES-mBlueberry2 vector also using BamHI digestion and verified by sequencing.

The constitutive lentiviral pRRL.SFFV.EGFP.miR-E vector was generated by subcloning the EGFP.miR-E from the pRRL LT3GECIR vector (generously provided by the Zuber lab) into the pRRL.SFFV.IRES.EGFP(6). Therefore the LT3GECIR vector was cut with MluI and self-ligated. The pRRL.SFFV.IRES.EGFP vector was cut with XhoI and EcoRI (Klenow) followed by self-ligation. The EGFP.miR-E was isolated from the LT3GECIR vector with BamHI and SacII digestion and cloned into the pRRL.SFFV.IRES.EGFP cut with BamHI and SacII, resulting into the pRRL.SFFV.EGFP.miR-E vector. Subsequently miR-E shRNAs for USP7 and a SCR control were designed or selected from Table S3 (Fellmann et al, 2013). USP7 #A: TGCTGTTGACAGTGAGCGACAGGATTTATTCAAGATACTATAGTGAAGCC ACAGATGTATAGTATCTTGAATAAATCCTGCTGCCTACTGCCTCGGA; USP7#B:TGCTGTTGACAGTGAGCGCC AAGAACATTGTTAAATTCAATAGTGAAGCCACAGATGTATTGAATTTAACAATGTTCTTGATGCCTACTGCCTC GGA; SCR: TGCTGTTGACAGTGAGCGCAGGAATTATAATGCTTATCTATAGTGAAGCCACAGATGTATAGATA AGCATTATAATTCCTATGCCTACTGCCTCGGA . 97-mer oligonucleotides were purchased from Invitrogen and PCR amplified using the primers miRE-Xho-fw (5’ TGAACTCGAGAAGGTATATTGCTGTTGACAGTGAGCG-3’) and miRE-EcoOligo-rev (5’-TCTCGAATTCTAGCCCCTTGAAGTCCGAGGCAGTAGGC-3’) according to Phusion High-Fidelity kit (Thermo Scientific). The PCR product (139 bp) was first ligated into pJet1.2 using Blunt-End cloning protocol and verified by sequencing. Then the miR-E shRNA was isolated using XhoI and EcoRI digestion and ligated into the pRRL.SFFV.EGFP.miRE vector also using XhoI and EcoRI digestion. Generation of lentiviral viruses and transductions were performed as described previously (7).

**Flow cytometry analysis**

Flow cytometry analyses were performed on the BD LSR II (Becton Dickinson (BD) Biosciences) and data were analysed using FlowJo (Tree Star Inc, Ashland, OR, USA). Cells were sorted on a MoFlo XDP or MoFLo Astrios (Beckman Coulter). For Hoechst/PyroninY staining, cells were resuspended in HPGM (Lonza, Leusden, The Netherlands) and stained with 5 ug/ml Hoechst 33342 (Invitrogen) at 37°C for 30-45 min. Then 1 ug/ml PyroninY (Sigma) was added and incubated for 30-45 min at 37°C. Upon FcR blocking (MACS miltenyi Biotec), cells were stained with CD34-APC (581, BD Biosciences) and CD38-AlexaFluor 700 (HIT2, Biolegend) at 4°C for 30 min. Cells were washed in medium containing Hoechst/PyroninY and analysed on the BD LSR II. *In vivo* engraftment levels were analysed in peripheral blood (PB), bone marrow, liver and spleen. Prior to staining, cells were blocked with anti-human FcR block (MACS miltenyi Biotec) and anti-mouse CD16/CD32 block (BD Biosciences) and stained with CD45-BV421 (HI30), CD19-BV785 (HIB19) and CD33-APC (WM53) all from Biolegend at 4°C for 30 min.

**Western blotting**

For detecting His-tagged ubiquitinated proteins cells were lysed in 1 ml denaturing lysis buffer (6M Guanidium-HCl, 100 mM $Na_2HPO_4$ x $2H_2O$, 10 mM Tris-HCl pH 8.0, 5 mM Imidazole, 10 mM β-mercaptoethanol) and sonicated on ice. Lysates were pre-cleared by centrifugation at 14000 rpm at 4°C and five volumes denaturing lysis buffer was added. His-tagged ubiquitinated proteins were purified by adding 75 µl pre-equilibrated Ni-NTA magnetic agarose beads (Jena Bioscience) and incubated for 4h on a rotating wheel. Beads were separated using a magnetic rack and washed 1x in denaturing lysis buffer without Imidazole, 1x in wash buffer pH 8.0 (8M Urea, 100 mM $Na_2HPO_4$ x $2H_2O$, 10 mM Tris-HCl pH 8.0 and 10 mM β-mercaptoethanol), 1x in wash buffer pH 6.3 (8M Urea,

100 mM $Na_2HPO_4$ x $2H_2O$, 10 mM Tris-HCl pH 6.3, 10 mM β-mercaptoethanol) plus 0.2% Triton X-100, and 1x in wash buffer pH 6.3 plus 0.1% Triton X-100. His-tagged ubiquitinated proteins were eluted from the beads by adding 75 µl elution buffer (200 mM Imidazole, 150 mM Tris-HCl pH 6.7, 30% glycerol, 5% SDS and 720 mM β-mercaptoethanol) and incubated for 20 min. on a rotating wheel. Elution samples were diluted 2x in Laemmli sample buffer containing 10% β-mercaptoethanol and immediately boiled for 5 min. prior to Western blot analysis.

GFP mediated pull out cell lysates, input, non-bound and bound fractions were boiled for 5-10 min. in Laemmli sample buffer prior to SDS-polyacrylamide gel electrophoresis. Proteins were transferred to fluorescence polyvinylidene difluoride (PVDF FL, Millipore) membrane by semidry blotting. Membranes were blocked in Odyssey blocking buffer (Westburg). The following primary antibodies were used: anti-USP7 (A300-033A, Bethyl Laboratories), anti-RING1B (ab 181140, Abcam), anti-GFP (sc-9996, Santa Cruz), anti-Ubiquitin (FK2, Enzo Life Sciences), anti-PCGF1 (ab183499, Abcam). Fluorescent secondary antibodies either goat anti-mouse IRDye 800 or goat anti-rabbit IgG (H+L) Alexa Fluor 680 (Invitrogen) were used for detection. Membranes were scanned using the Odyssey CLx Imaging System (Li-Cor Biosciences).

**RNA seq analysis and quantitative real-time PCR**

RNA samples for sequencing were prepared for DMSO and P22077 (30 µM) treated K562 cells at 4h, 8h, 16h and 24h. Total RNA was isolated using the RNeasy Mini Kit from Qiagen (Venlo, The Netherlands) according to the manufacturer’s recommendations. Initial quality check and RNA quantification of the samples was performed by capillary electrophoresis using the LabChip GX (Perkin Elmer). Sequence libraries were generated with 50 ng mRNA, using Lexogen Quantseq 3’ prep kit (Lexogen GmbH) according to the manufacturer’s recommendations. The obtained cDNA fragment libraries were sequenced on an Illumina NextSeq500 using default parameters (single read). Bioinformatics were performed on the Strand Avadis NGS (v3.0) software (Strand Life Sciences Pvt.Ltd). Sequence quality was checked for GC content, base quality and composition using FASTQC and StrandNGS. Quality trimmed reads were aligned to build Human Hg19 transcriptome. Ensembl Genes and transcripts (2014.01.02) was used as gene annotation database. Quantified reads were normalized using the DESeq package. Reads with failed vendor QC, quality score less than 24 (average), mapping quality score below 50 and length less than 20 were all filtered out.

For quantitative RT-PCR, RNA was reverse transcribed using the iScript cDNA synthesis kit (Bio-Rad) and amplified using SsoAdvanced SYBR Green Supermix (Bio-Rad) on a CFX384 Touch Real-Time PCR Detection System (Bio-Rad). RPL27 was used as housekeeping gene. Primer sequences are available on request.

**USP7 inhibition *in vivo***

Eight to ten week old female NSG (NOD.Cg-Prkdcscid Il2rgtm1Wjl/SzJ) mice were purchased from the Centrale Dienst Proefdieren (CDP) breeding facility within the University Medical Center Groningen. Mouse experiments were performed in accordance with national and institutional guidelines, and all experiments were approved by the Institutional Animal Care and Use Committee of the University of Groningen (IACUC-RuG). 24h prior to transplantations, mice were sub-lethally irradiated with a dose of 1.0 Gy (X-RAD 320 Unit, PXINC 2010). After irradiation mice received Neomycin (3.5 g/l) in their drinking water and soft food (RM Convalescence + BG SY (M); Special Diet Services; Witham, England) for two weeks. For secondary transplantations, 5 x $10^4$ MLL-AF9 EGFP cells from primary leukemic mice (CB MLL-AF9 xenograft mouse model, (1, 8)) were injected IV (lateral tail vein). Peripheral blood chimerism levels were monitored by regular blood sample analysis. Mice were randomly divided into two groups, weighted and treated with DMSO as control (n=5) or 20 mg/kg P22077 (n=6) via intraperitoneal (IP) injections daily starting four weeks post-transplant. Prior to injections, P22077 was dissolved in DMSO (or DMSO only as control) and directly mixed with Cremophor EL (1:1). This solution was then diluted 1:4 in saline, to get an end concentration of max. 10% DMSO. Mice were humanely terminated by cervical dislocation under isoflurane anesthesia when chimerism levels in the blood exceeded 40%. Peripheral blood, bone marrow, spleen and liver were analysed.
